# Molecular Dynamics Simulations of HLA-CW4-B2M-KIR2DL1 Protein and Homology Modeling of a Complex Associated with Psoriasis Disease (HLA-CW6-B2M-KIR2DS1)

**DOI:** 10.1101/2021.12.20.473468

**Authors:** Mansour H. Almatarneh, Ahmad M. Alqaisi, Enas K. Ibrahim, Ghada G. Kayed, Joshua W. Hollett

## Abstract

Molecular dynamics (MD) simulation was used to study the interactions of two immune proteins of HLA-Cw4-β2m-KIR2DL1 complex with small peptide QYDDAVYKL (nine amino acids) in an aqueous solution. This study aims to gain a detailed information about the conformational changes and the dynamics of the complex. The right parameters and force field for performing the MD simulations that was needed to calibrate the complex structure were determined. The non-bonded interactions (Electrostatic and van der Waals contributions), H-bond formation, and salt bridges between the ligand HLA-Cw4 and the receptor KIR2DL1 were estimated using the obtained MD trajectories. The buried surface area due to binding was calculated to get insight into the causes of specificity of receptor to ligand and explains mutations experiment. The study concluded that β2-microglobulin, one part of the complex, is not directly interacting with the peptide at the groove; therefore, it could be neglected from simulation. Our results showed that β2-microglobulin does not have any significant effect on the dynamics of the 3D-structure of the complex. This project will help in understanding to optimize candidate drug design, a small peptide that disrupts the interaction, for the optimal biological effect.

## 1. Introduction

Major Histocompatibility Complex (MHC) refers to a group of genes that encodes for somatic cell-surface proteins in immunology. The surface proteins play a crucial role in the immune response by presenting T cells with peptides. This group includes two subfamilies, class I and class II MHC, the two classes of families I and II are usually linked together in a single gene complex[1]. In humans, this single gene complex is located on chromosome 6, and it is called Human Leukocyte Antigen (HLA). Proteins expressed on the surface of all nucleated somatic cells are polymorphic class I, MHC molecules (Fig. 1). Class, I human MHC-HLA antigens mediate response by natural killer (NK) cells. They work to present cytotoxic T lymphocytes (CTL) with peptides and consist of the following two chains: An alpha or heavy chain consists of three extracellular domains (α1, α2, and α3), a transmembrane region, a cytoplasmic domain, and β2-microglobulin (*β2m*), Consisting of a single domain.

**Fig. 1.**
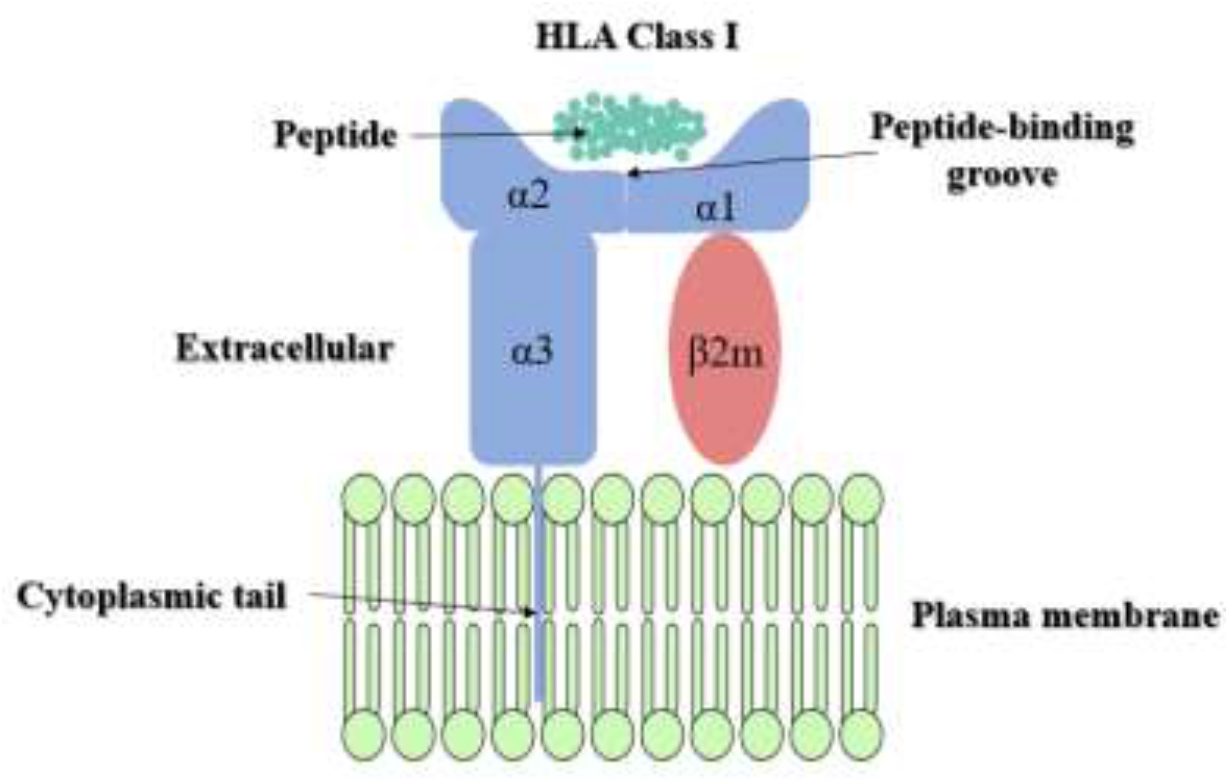
MHC class I molecules.

The gain from this project will help us to construct the protein complex that is associated with psoriasis disease by superposition and homology modeling that based on the HLA-Cw4-β2m-KIR2DL1-QYDDAVYKL complex. Psoriasis is a chronic inflammatory disease of the skin[2–4]. This disease comes with different variants of clinical subtypes and mainly affects the skin and joints, causing red scaly patches. Psoriasis can develop anywhere on the human body, but most commonly on the trunk, elbows, knees, and scalp[5]. Psoriasis’s significant genetic origins were supported by the association of the disease with Human Leukocyte Antigen (HLA), particularly HLA-Cw6 allele type[6, 7].

Molecular Dynamics (MD) simulations are one of the principal tools in the computational study of biological molecules. It is used to investigate the structure, dynamics, and thermodynamics of biological molecules and their complexes[8–10]. In this work, using MD simulations, we study the interactions between the HLA-Cw4 Antigen, β2-macroglobulin (β2m), and HLA-Cw4 specific peptide (one-letter code amino acid sequence (QYDDAVYKL))[11]. These are bound to HLA natural killer cell inhibitory receptor KIR2DL1. This complex will be referred to in this paper as HLA-Cw4-β2m-KIR2DL1. This complex structure will give us an excellent model to study the HLA-Cw6-β2m-KIR2DS1 complex associated with many aspects of psoriasis disease[6].

The parameters for running the MD simulations, box dimensions, the right force field, and the time (in *ns*) needed to calibrate the complex structure were determined. From there, the model of the complex of HLA-Cw6, β2-microglobulin (β2m), and KIR2DS1 will be constructed by superposition and homology modeling using SWIS-MODEL based on the HLA-Cw4-β2m-KIR2DL1 complex. The future optimal goal is to provide a detailed understanding of the HLA-Cw6-β2m-KIR2DS1, which is not yet known, which in turn will benefit health science and drug design in developing more effective treatments for psoriasis and other inflammatory diseases. This computational study will benefit experimentalists by providing detailed study about this complex, aid in the design of new experiments for further understanding and help develop synthetic procedures for this complex. This knowledge will also benefit health science and drug design

## 2. Computational Methods

The initial HLA-Cw4-*β*2m-KIR2DL1 structure, composed of 4 chains, was retrieved from the protein databank [PDB entry ***lim9***] determined at 2.70 *Å* resolution[12]. All the simulations were performed using the NAMD 2.14 package [13, 14] and the CHARMM36 force field[15]. The protein was solvated using the explicit TIP3P solvation model[16] and the periodic boundary conditions were used with a dimension of 15.2 *nm*^*3*^. The Isobaric-isothermal (NPT) ensemble and a 2 *fs* time step of integration were chosen for all MD simulations. The MD protocols involved minimization, annealing, equilibration, and production, with minimization of 4 *ps*, annealing processes for 288 *ps*, equilibration of 1 *ns*, and MD simulation production of 100 *ns*. The temperature was set to 300.15 *K* using the Langevin thermostat[17]. Long-range Coulombic interactions were treated using Particle-mesh Ewald (PME) method[18, 19]. All covalent bonds involving hydrogen atoms were constrained using the SHAKE algorithm[20] and Leonard-Jones interactions were treated with a switching feature that limits the cut-off distance to 18 Å.

Homology modelling (comparative modelling) is one method of predicting the computational structure used to determine the 3D-protein structure from its amino acid sequence. It is considered the most reliable tool for predicting the computational structure[21]. To create 3D models for the target sequence based on its templates, different methods are used. It is possible to identify model building approaches as rigid body-assembly techniques, section matching techniques, methods of spatial constraint, and methods of artificial evolution[22]. The protein structure is broken down into simply retained core regions, loops and side chains in the rigid-body assembly. This approach depends on the natural dissection that enables creating a 3D protein structure by putting together these rigid bodies that are collected from the aligned structure[23]. There are many tools can do this, like SWISS-MODEL[24].

## 3. Results and discussion

The 100 *ns* MD trajectories obtained for the complex HLA-Cw4-β2m-KIR2DL1 with the peptide were produced using NAMD. It was analyzed using the visual molecular dynamics software (VMD)[25] to gain more detailed information about the structure, thermodynamics, and dynamics of intermolecular interactions between the HLA-Cw4 gene product and its receptor, KIR2DL1. The protocol of MD simulations of the HLA-Cw4 will help to optimize the best parameters for the calculation needed for future work on studying HLA-Cw6-β2m-KIR2DS1 protein complex. Analysis of the molecular dynamics (MD) simulations of the immune protein HLA-Cw4 is discussed below, including comparing the theoretical results with experimental findings.

### 3.1. Structural Stability

The X-ray structure of the HLA-Cw4 complex was run for 100 *ns* where energies were recorded every 200 *ps* and coordinates every 2000 *ps* to achieve equilibrium as shown in Fig. 2. The assessment of the structural stability for the MD simulation was achieved by the root mean squared deviation (RMSD) and root mean squared fluctuation (RMSF). The average value of the RMSD for the protein backbone over the 100 *ns* MD simulation was of 2.91 Å. Thus, a simulation time of 100 *ns* is long enough to achieve equilibrium. Notably, in the last 20 *ns* of simulation, the RMSD value is less than 2.9 Å. Therefore, the average values of nonbonded interactions for the last 20 *ns* were calculated and discussed. It should be mentioned that the value of RMSD for the conserved residues of the active site about 2-3 Å.

**Fig. 2.**
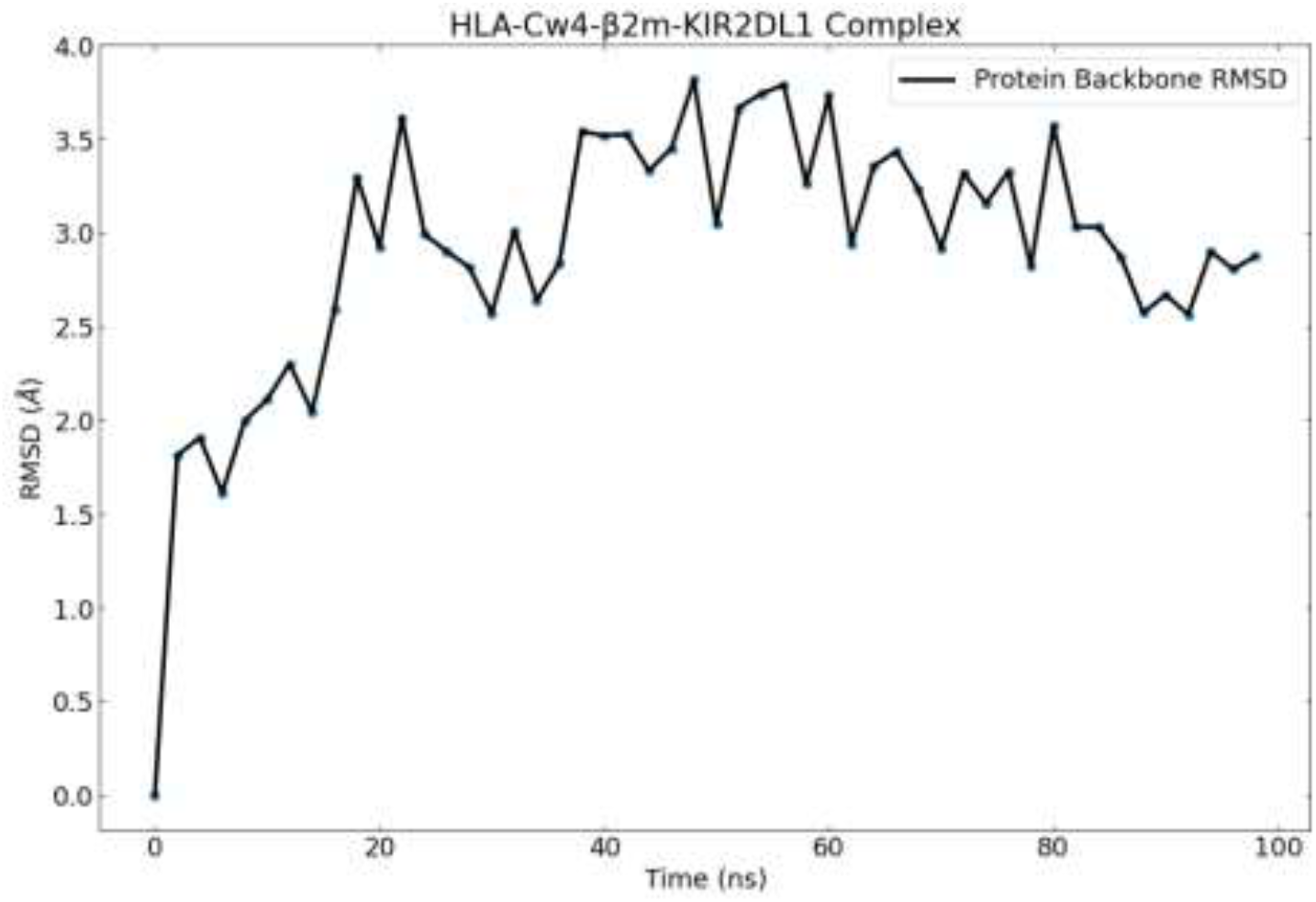
RMSD versus MD simulation time for the HLA-Cw4-β2m-KIR2DL1 protein backbone The RMSF values that represent the average over time of the RMSD over protein backbone atoms. The average value of the RMSF for the protein backbone was 0.484 Å, as shown in Fig. 3.

**Fig. 3.**
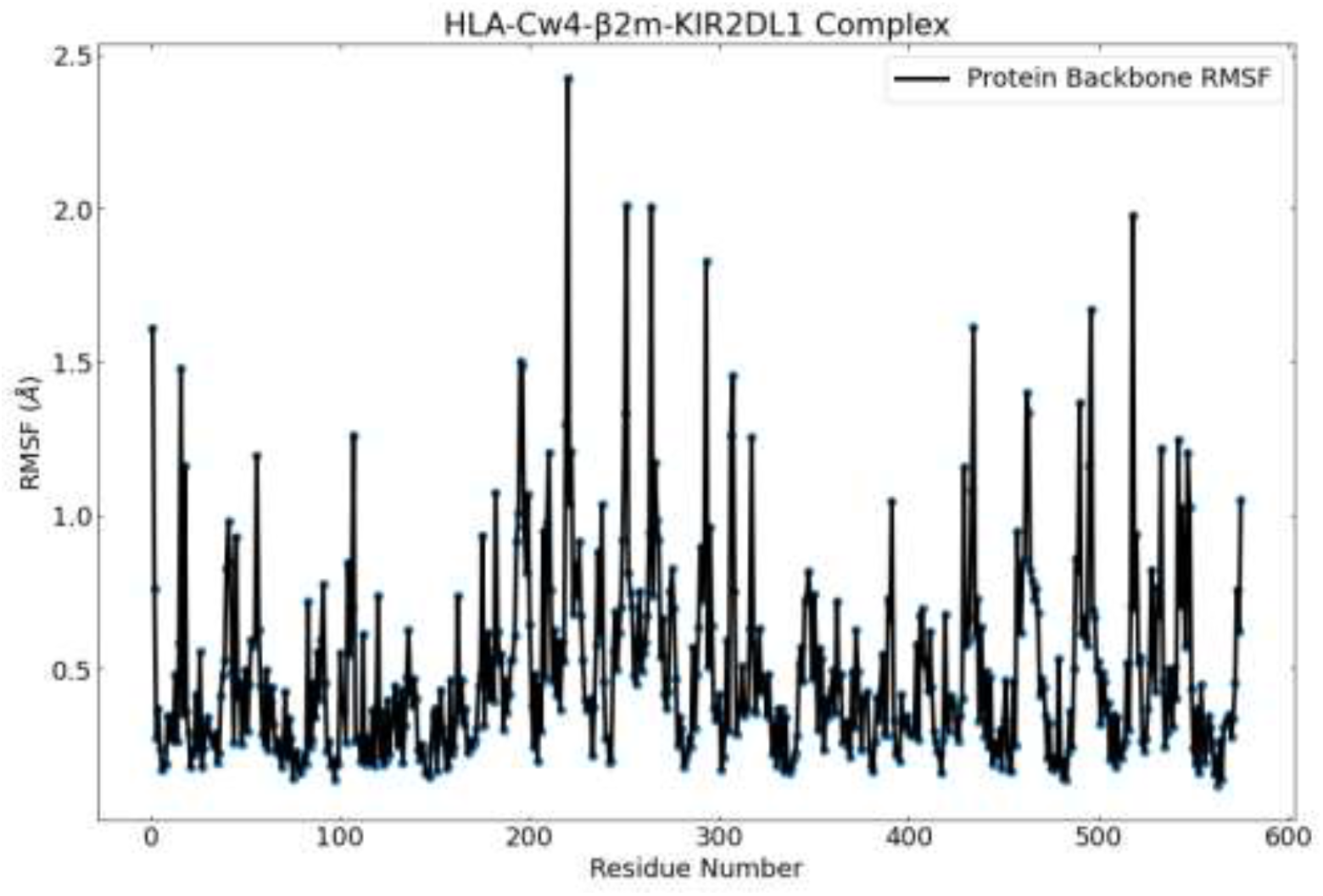
RMSF versus MD simulation time for the HLA-Cw4-β2m-KIR2DL1 protein backbone.

Ramachandran plot analysis indicates whether the backbone torsion angles *(φ/Ψ)* of our structure are within the allowed region. Most of the peptide residues have *(φ/Ψ)* angles typical for an extended β strand and α helix. On the other hands, some residues adopt *(φ/Ψ)* angles that are found in a left-handed helix, and they are usually observed in residues forming tight turns and kinks such as Asp^57^, Asp^220^, Asn^42^, Asn^86^, Leu^130^, Arg^151^ and others due to the formation of internal hydrogen bonds in the chain. These observations agree with the previously reported *X*-ray study for HLA-Cw4 and KIR2D complex[11]. However, some points appear in the sterically disallowed region due to Gly amino acids (Fig. 4).

**Fig. 4.**
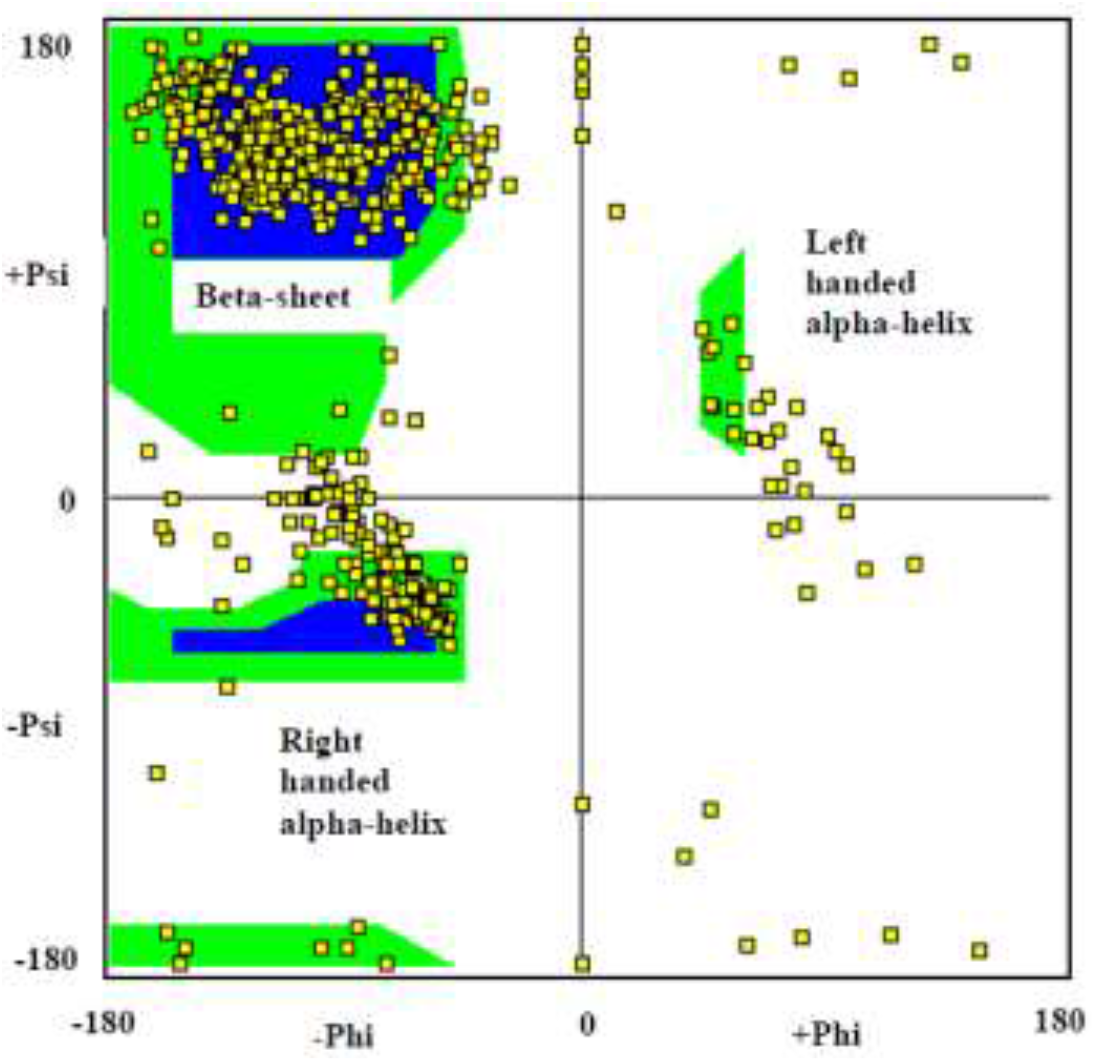
Ramachandran Plot for the final frame of the 100 *ns* MD simulation.

### 3.2. Hydrogen Bonding

Hydrogen bonding numbers and occupancies were calculated using *HBonds Plugin*, version 1.2 in VMD software. Hydrogen bond occupancy is computed as the fraction of conformations out of 50 conformations of the complex system in which the given residue participates in a hydrogen bond. The 50 conformations of the complex were derived from the corresponding 100 *ns* molecular dynamics trajectory. The hydrogen bonding cut-off distance was set to 3.4 Å. Figs. 5 and 6 depict the hydrogen bonding between the HLA-Cw4 and the KIR2DL1 interface and the KIR2DL1 interface, respectively. We find several hydrogen bonds between the ligand HLA-Cw4 (chain A) and receptor KIR2DL1 (chain D), such as: (Lys^146^, Asp^183^), (Tyr^84^, Asp^183^), (Ala^149^, Glu^106^), (Lys^80^, Glu^187^), Arg^145^ with each of (Asp^145^, Ser^133^). That is in agreement with a previous study[12], H-bond formation between Lys^146^ of the HLA-Cw4 and Glu^187^ of the KIR2DL1 contributes to the specificity of receptor to ligand (occupancy = 30.00%), which was previously reported[12], see Fig. 5. Besides, other new hydrogen bonds appear in our simulation between ligand and receptor such as: (Arg^151^, Glu^106^), (Arg^75^, Asp^47^), and (Arg^79^, Met^44^). Furthermore, in our simulation, we detect new hydrogen bond with high occupancy (36.00%) between Lys^80^ of HLA-Cw4 and Asp^183^ of KIR2DL1, which it is expected to contribute to specificity of receptor to ligand too. Table 1 summarizes the hydrogen bonds between the ligand HLA-Cw4 and the receptor KIR2DL1.

**Table 1.** Hydrogen bonds at the KIR2DL1 and HLA-Cw4 interface.

| HLA-Cw4 | KIR2DL1 | Occupancy |
| --- | --- | --- |
| Arg<sup>75</sup> - Side - NH2 | Asp<sup>47</sup> - Side - OD2 | 68.00% |
| Arg<sup>75</sup> - Side - NH1 | Asp<sup>47</sup> - Side - OD1 | 54.00% |
| Ala<sup>149</sup> - Main - O | Glu<sup>106</sup> - Main - N | 50.00% |
| Tyr<sup>84</sup> - Side - OH | Asp<sup>183</sup> - Side - OD2 | 42.00% |
| Lys<sup>146</sup> - Side - NZ | Asp<sup>183</sup> - Side - OD1 | 36.00% |
| Arg<sup>151</sup> - Side - NH2 | Glu<sup>106</sup>-Side-OE2 | 24.00% |
| Lys<sup>80</sup> - Side - NZ | Glu<sup>187</sup> - Side - OE2 | 22.00% |
| Arg<sup>151</sup> - Side - NH1 | Glu<sup>106</sup> - Side - OE2 | 22.00% |
| Arg<sup>151</sup> - Side - NH2 | Glu<sup>106</sup> - Side - OE1 | 20.00% |
| Arg<sup>79</sup> - Side - NE | Met<sup>44</sup> - Main - O | 20.00% |
| Arg<sup>151</sup> - Side - NE | Glu<sup>106</sup> - Side - OE1 | 18.00% |
| Lys<sup>80</sup> - Side - NZ | Asp<sup>183</sup> - Side - OD2 | 16.00% |
| Arg<sup>145</sup> - Side - NE | Ser<sup>133</sup> - Side - OG | 16.00% |
| Arg<sup>145</sup> - Side - NH2 | Ser<sup>133</sup> - Main - O | 16.00% |
| Arg<sup>145</sup> - Side - NH2 | Asp<sup>135</sup> - Side - OD1 | 14.00% |
| Arg<sup>151</sup> - Side - NH1 | Glu<sup>106</sup> - Side - OE1 | 12.00% |
| Tyr<sup>84</sup> - Side - OH | Asp<sup>183</sup> - Side - OD1 | 8.00% |
| Lys<sup>146</sup> - Side - NZ | Glu<sup>187</sup> - Side - OE2 | 8.00% |
| Lys<sup>80</sup> - Side - NZ | Glu<sup>187</sup> - Side - OE1 | 8.00% |

**Fig. 5.**
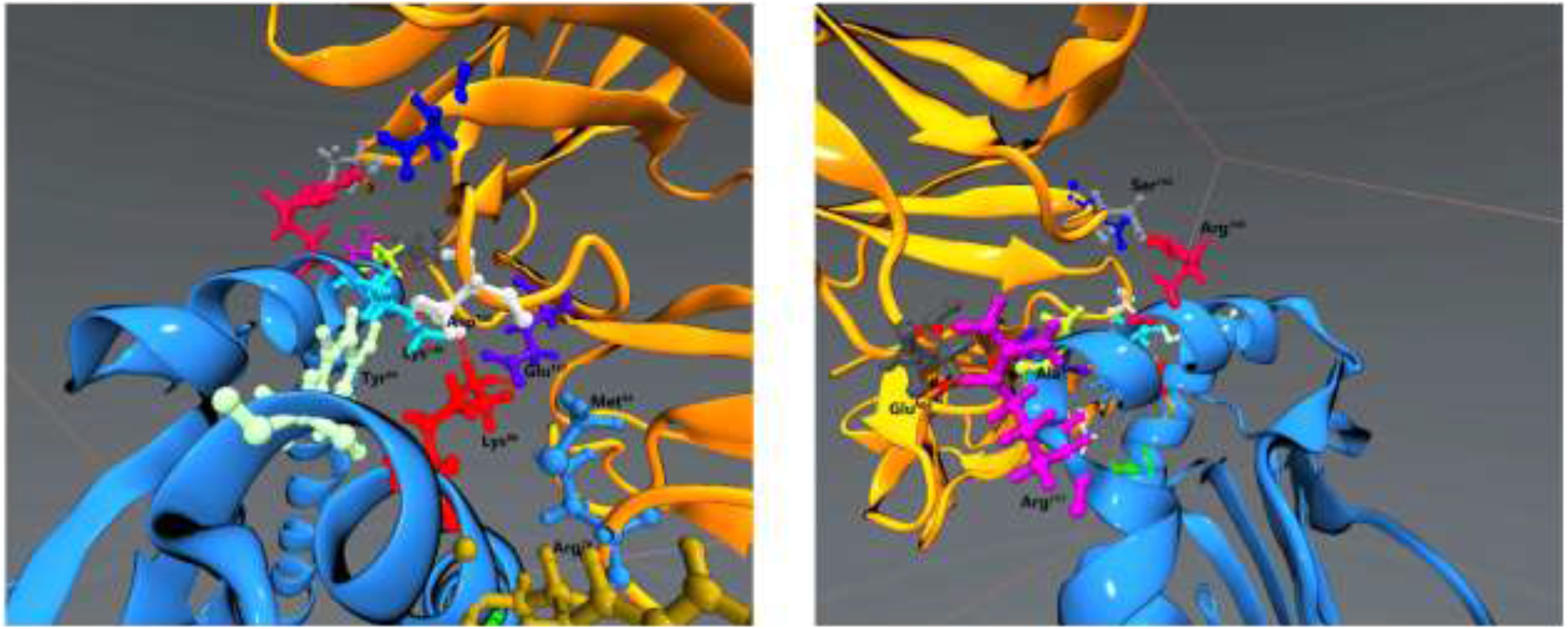
Hydrogen Bonding at the HLA-Cw4 and the KIR2DL1 interface.

**Fig. 6.**
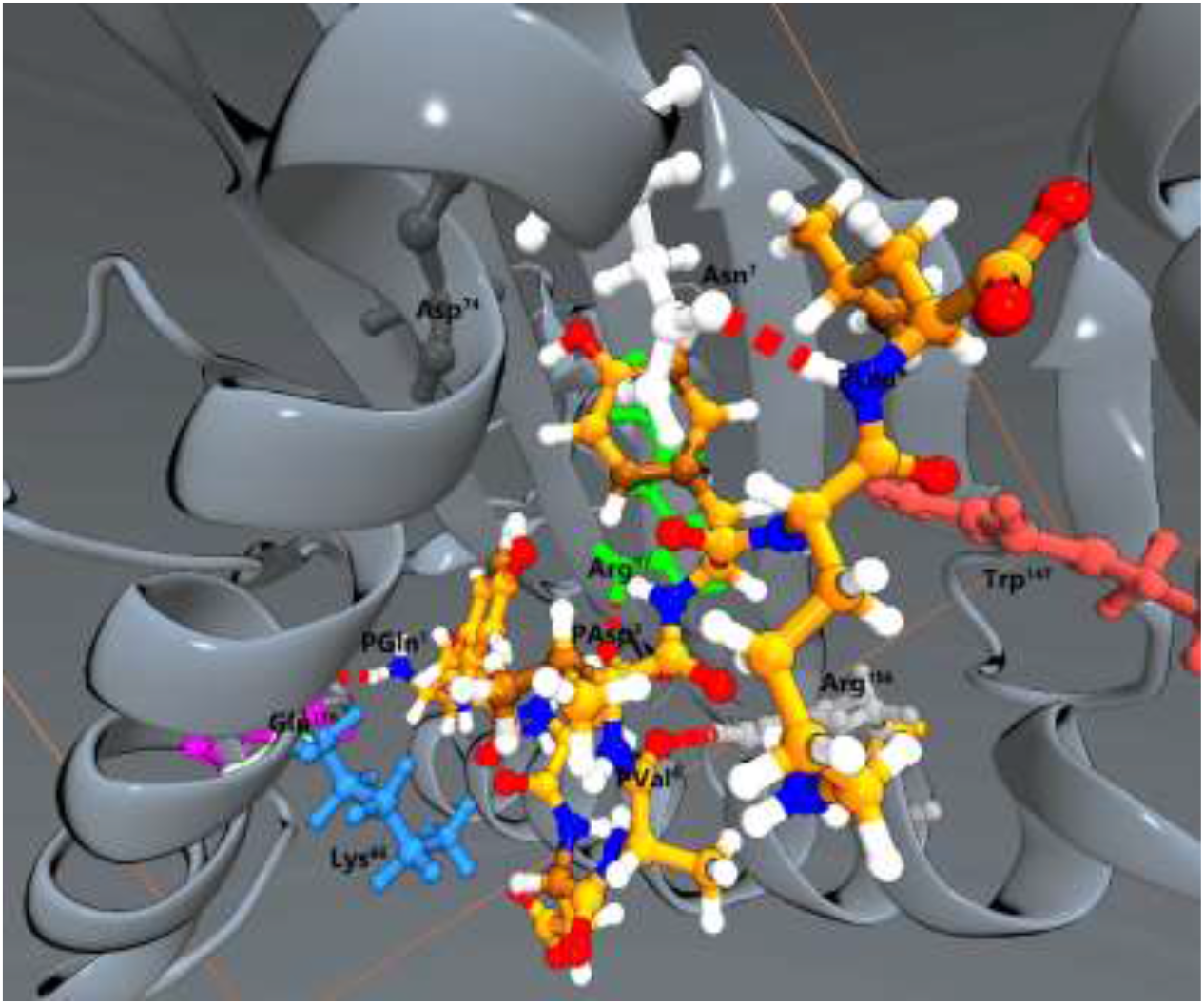
Hydrogen Bonding between HLA-Cw4 and HLA-Cw4 specific peptide (QYDDAVYKL).

We also find the extensive hydrogen bonding pattern between the ligand HLA-Cw4 and the HLA-Cw4 specific peptide, which is described previously in the literature[11] (Fig. 6). Figs. S1-S5 in the Supplementary Information (SI) depict inter-molecular H-bonds between HLA-Cw4 (chain A) and KIR2DL1 (chain D). Table S1 in the SI summarizes the hydrogen bonds between the ligand HLA-Cw4 and its specific peptide (QYDDAVYKL).

### 3.3. Salt Bridge Interactions

The salt-bridge interactions in protein are formed between oppositely charged amino acids, which cause electrostatic interactions. They contribute to the specificity of receptor to ligand[26]. According to Table 2, several salt bridges formed between the residues in receptor KIR2DL1 (Chain D) and ligand HLA-Cw4 (Chain A). The salt-bridge between Lys^80^ of HLA-Cw4 and GLu^187^ of KIR2DL1 contribute to the specificity of receptor to the ligand. They are previously reported in x-ray analysis of the complex of HLA-Cw4-KIR2DL1[12].

**Table 2.** Salt bridge between receptor (Chain D) and ligand (Chain A).

| Residue | Chain | Residue | Chain | Avg No. of salt bridge for the last 20 ns |
| --- | --- | --- | --- | --- |
| Asp<sup>183</sup> | Chain D | Lys<sup>146</sup> | Chain A | 2.90351 |
| Glu<sup>21</sup> | Chain D | Arg<sup>69</sup> | Chain A | 8.0569 |
| Glu<sup>187</sup> | Chain D | Lys<sup>80</sup> | Chain A | 4.2425 |
| Asp<sup>135</sup> | Chain D | Arg<sup>145</sup> | Chain A | 3.7906 |
| Asp<sup>183</sup> | Chain D | Lys<sup>80</sup> | Chain A | 3.4609 |
| Asp<sup>72</sup> | Chain D | Arg<sup>69</sup> | Chain A | 5.0498 |
| Glu<sup>187</sup> | Chain D | Lys<sup>146</sup> | Chain A | 4.6718 |
| Asp<sup>47</sup> | Chain D | Arg<sup>75</sup> | Chain A | 7.1159 |

### 3.4. Nonbonded interactions in water

It is crucial to investigate the nonbonded interactions between the ligand, the HLA-Cw4, and the receptor KIR2DL1 in water. If there are enough interactions between the ligand and receptor, the ligand HLA-Cw4 of normal cells succeeds in switching off the natural killer cell by binding to the receptor KIR2DL1. On the other hand, if there are no enough nonbonded interactions between ligand and receptor due to mutant in specific amino acids, the ligand will not bind to the receptor. Therefore, in this case, the natural killer cells attack the normal cell, which causes the illness. In the following sections, the nonbonded interactions between the chains and certain amino acids of ligand and receptor were discussed. Therefore, we can detect the cause of non-bonded interactions between ligand and receptor, which is responsible for eliminating the illness.[12] As you can notice in Fig. 7, the main source of non-bonded interaction between the ligand and receptor is due to interactions between chain A of ligand and chain D of the receptor, and it is mainly due to electrostatic interactions and minor vdW interactions. That is consistent with the previous study[12], which states that KIR2DL1 interacts with complementary charged residue for HLA-Cw4-β2m. The electropositive binding surface of HLA-Cw4-β2m interacts with the electronegative binding surface of KIR2DL1.

**Fig. 7.**
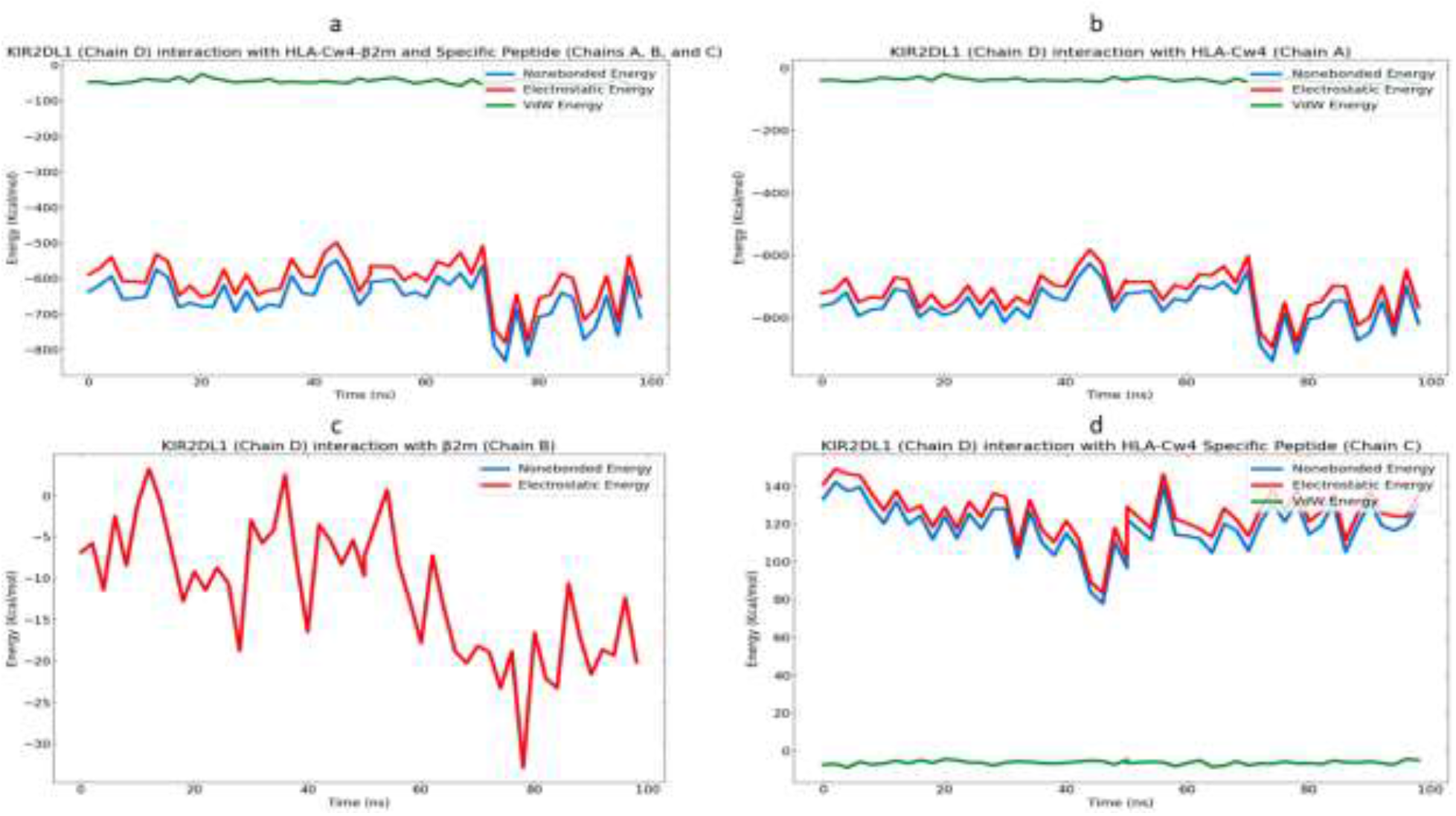
Non-bonded interaction energies of KIR2DL1 in water with (a): HLA-Cw4-β2m and HLA-Cw4 Specific peptide (b): HLA-Cw4 (c): β2m, and (d): HLA-Cw4 Specific peptide, estimated over a period of 100 *ns*.

According to Fig. 7, the non-bonded interactions between the chain B and receptor are less than chain A. And almost there are no non-bonding interactions between chain C and the receptor. As you can notice clearly, the interaction of chain C (small peptide) does not contribute to the binding of the ligand to the receptor. That is in agreement with the previous study[12], which states that peptide does not participate in recognition of HLA-Cw4 by KIR2DL1.

According to our results (Table S2 in SI), the main non-bonded interactions is due to the electrostatic interactions between the following amino acids residues in the receptor and the ligand, respectively: (Asp^183^, Lys^146^), (Glu^106^, Arg^151^), (Asp^135^, Arg^145^), (Glu^187^, Lys^80^) and (Ser^133^, Arg^145^). That is in consistent with the previous study for the main polar interactions between key amino acids for the ligand and the receptor.[12] In addition, the specificity of the receptor to the ligand is attributed to the interaction between Lys^80^ of HLA-Cw4 and Glu^187^ of KIR2DL1 (Fig. 8).[12] In agreement with our finding, the non-bonding interactions are relatively large about −903.7 kJ/mol and mainly electrostatic interaction. However, there are no polar interactions between the pairs (Met^44^, Lys^80^) and (Ser^184^, Lys^146^), which are mentioned in the previous study.[12]

**Fig. 8.**
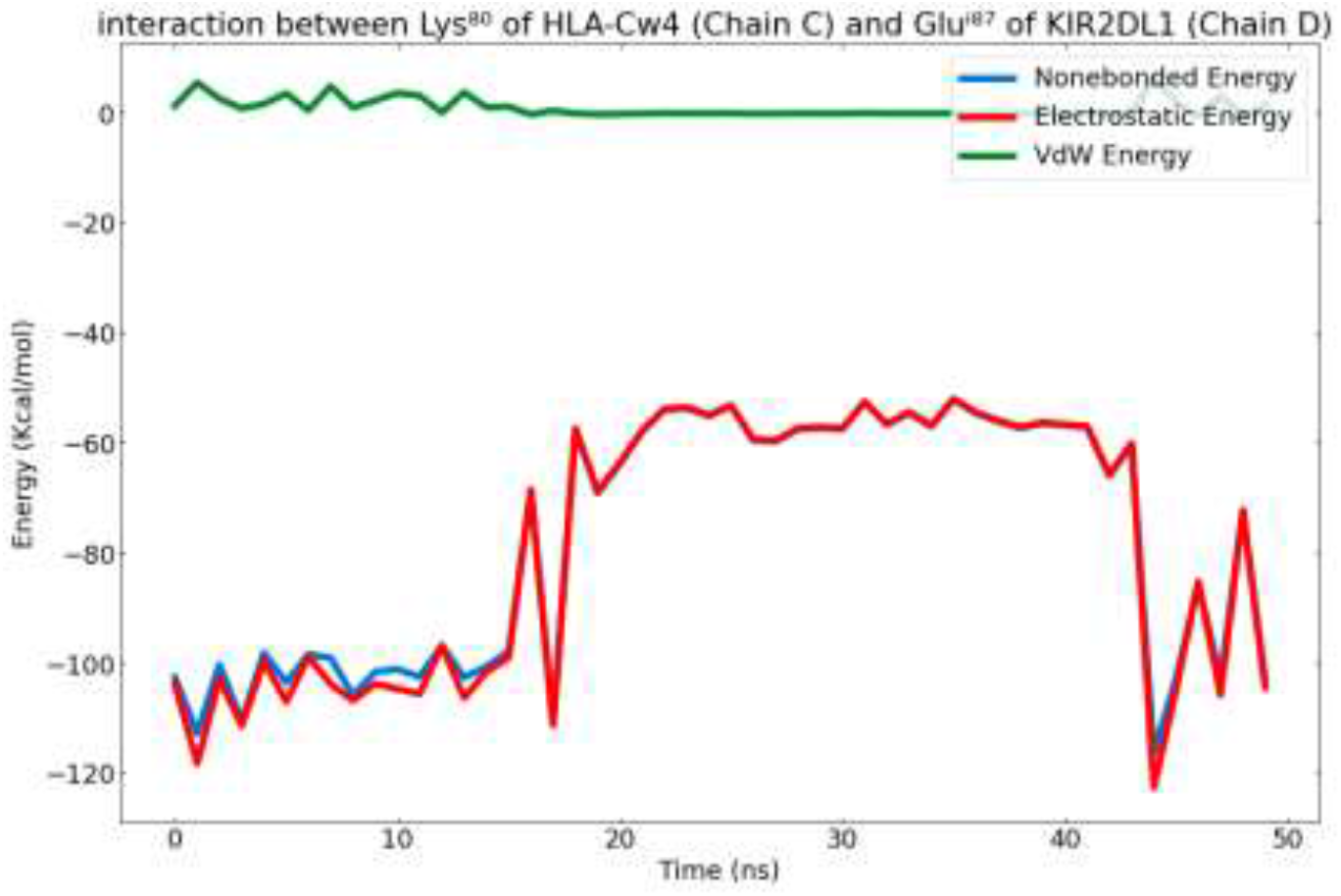
Non-bonded interaction energies between Lys^80^ of HLA-Cw4 and Glu^187^ of KIR2DL1 in water, estimated over a period of 100 *ns*.

Besides, there are *vdW* interactions among some amino acid residues of receptor and ligand. For example, (Met^44^, Arg^79^), (Tyr^105^, Ala^149^), (Phe^45^, Arg^75^), (Tyr^105^, Lys^146^), (Asp^72^, Val^76^), (Phe^181^, Lys^146^), (Phe^45^, Va^l76^) (Phe^45^, Arg^79^) and (Met^44^, Lys^80^). That is in agreement with the previous study for hydrophobic contact.[12] As we can notice, the *vdW* contribution to non-bonded interactions is less than electrostatic interactions. According to the previous studies[12] presence of a mutant in Lys^80^ of HLA-Cw4 causes KIR2DL1 not to recognize the HLA-Cw4. Table 3 reports the polar interactions and hydrophobic contacts between the HLA-Cw4 and the receptor KIR2DL1. According to our results, the *vdW* interactions are the leading non-bonding-interactions between Met^44^ of the receptor and Lys^80^ of the ligand, about −11.3 KJ/mol.

**Table 3.** Polar interactions and hydrophobic contacts at the between the HLA-Cw4 and the receptor KIR2DL1.

| Polar Interactions |  | Hydrophobic contacts |  |
| --- | --- | --- | --- |
| KIR2DL1 | HLA-CW4 | KIR2DL1 | HLA-Cw4 |
| Met<sup>44</sup> | Arg<sup>79</sup> | Met<sup>44</sup> | Arg<sup>79</sup>, Lys<sup>80</sup> |
| Glu<sup>106</sup> | Arg<sup>151</sup> | Phe<sup>45</sup> | Arg<sup>75</sup>, Val<sup>76</sup>, Arg<sup>79</sup> |
| Ser<sup>133</sup> | Arg<sup>145</sup> | Met<sup>44</sup> | Lys<sup>80</sup> |
| Asp<sup>135</sup> | Arg<sup>145</sup> | Tyr<sup>105</sup> | Ala<sup>149</sup> |
| Asp<sup>183</sup> | Lys<sup>146</sup> | Ser<sup>132</sup> | Ala<sup>149</sup> |
| Tyr<sup>105</sup> | Lys<sup>146</sup> | Ser<sup>133</sup> | Arg<sup>145</sup> |
| Glu<sup>187</sup> | Lys<sup>80</sup> | Phe<sup>181</sup> | Lys<sup>146</sup> |
| Asp<sup>72</sup> | Val<sup>176</sup> | Ser<sup>184</sup> | Lys<sup>146</sup> |
| Asp<sup>183</sup> | Tyr<sup>84</sup> | Glu<sup>187</sup> | Lys<sup>80</sup> |

*Contact Area and Solvent Accessible Surface Area*

According to a previous study by Fan and co-authors[11], Lys^80^ of HLA-CW4 is highly exposed to the receptor (KIR2DL1) (68% of its surface area exposed). Fig.9 displays the contact area between the Lys^80^ of HLA-CW4 and receptor (KIR2DL1). According to the contact area plot, the average area of contact is 86.3 A^2^, in agreement with our results. This may explain the role of Lys^80^ of the HLA-Cw4 in recognition of receptor (KIR2DL1) to the ligand HLA-Cw4. The experimental mutational experiment shows that Lys^80^ of ligand controls the specificity of the receptor to the ligand.[27]

**Fig. 9.**
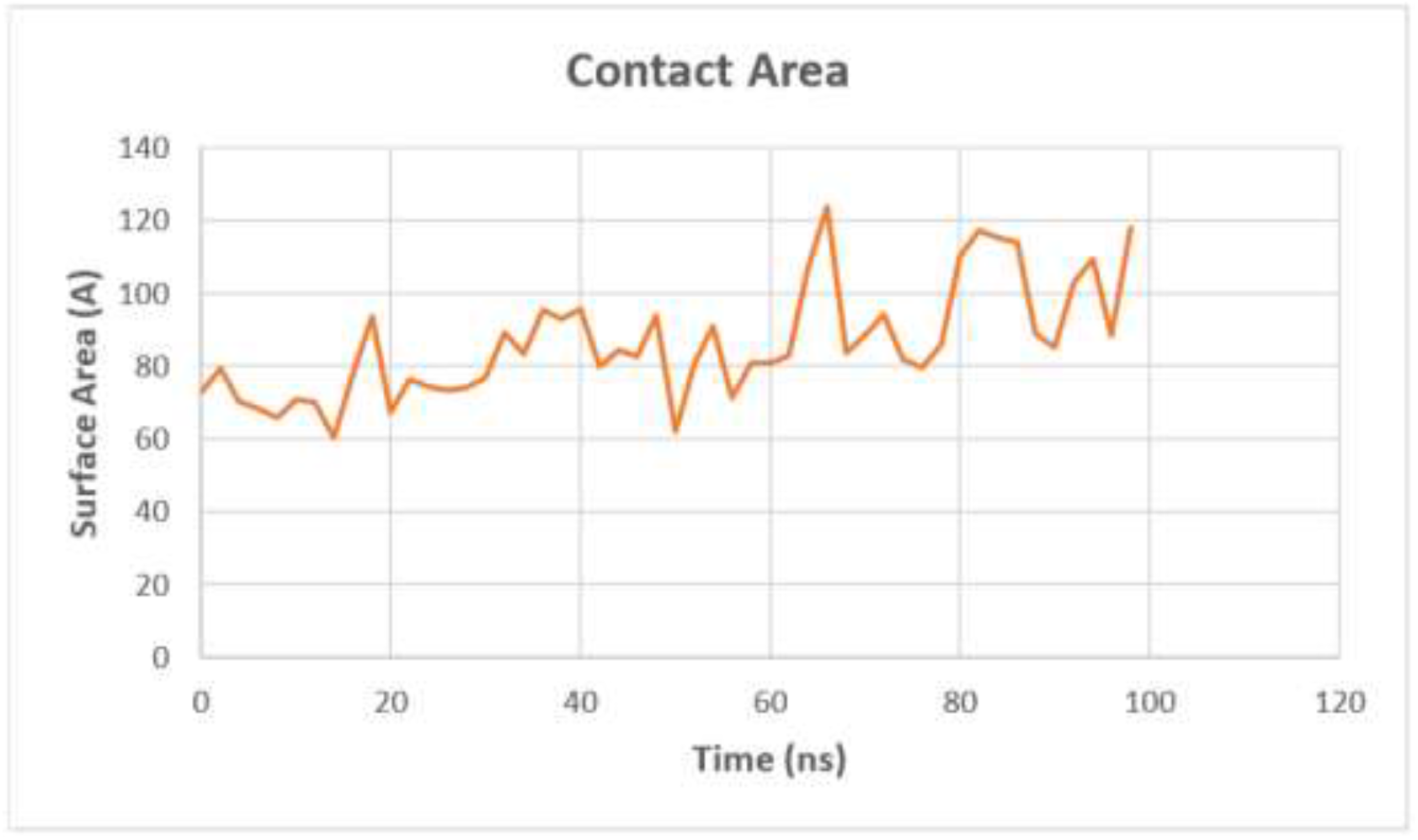
The contact area between the Lys80 of HLA-Cw4 and receptor (KIR2DL1).

**Fig. 10.**
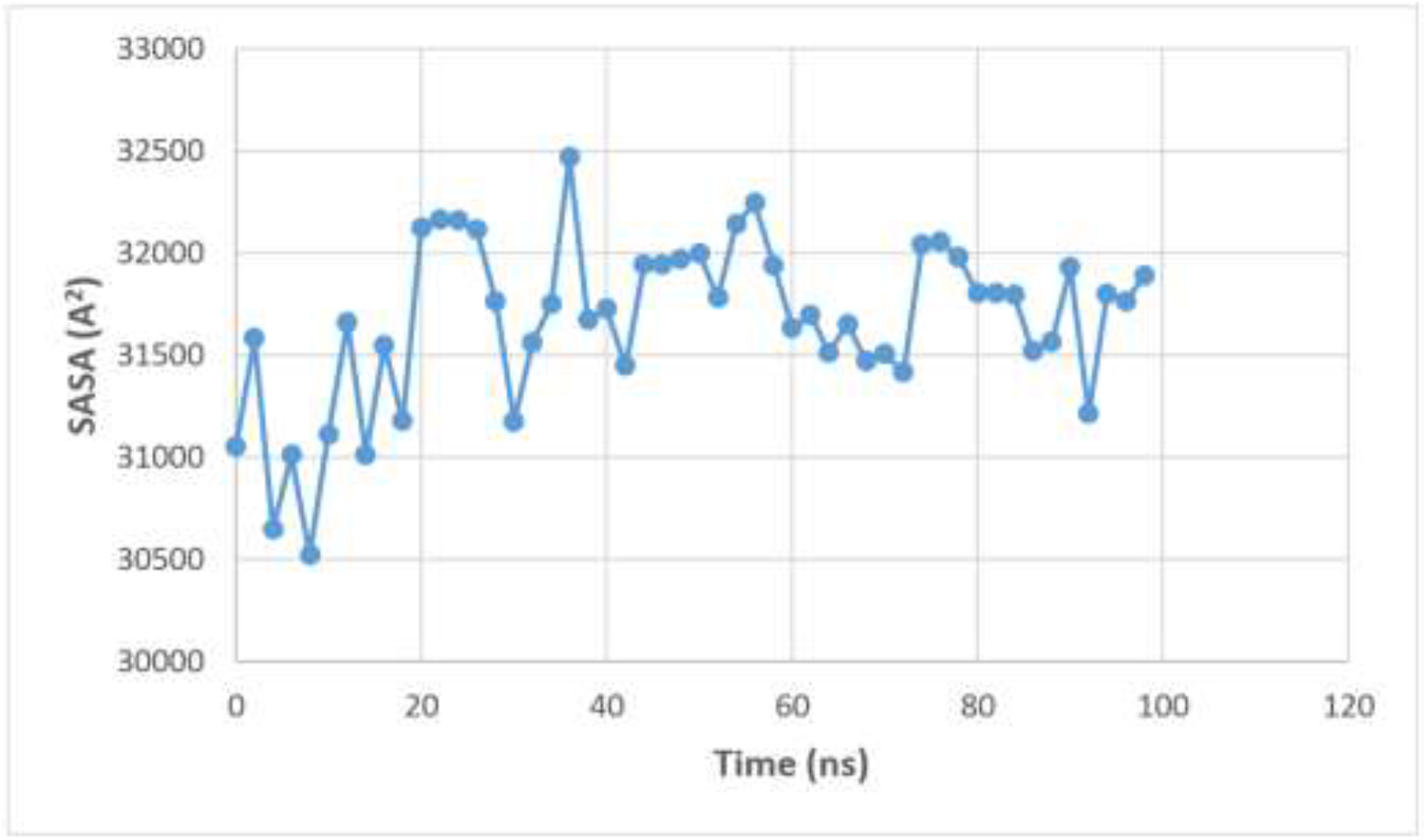
Solvent accessible surface area for HLA-Cw4-β2m-KIR2DL1 complex.

**Fig. 11.**
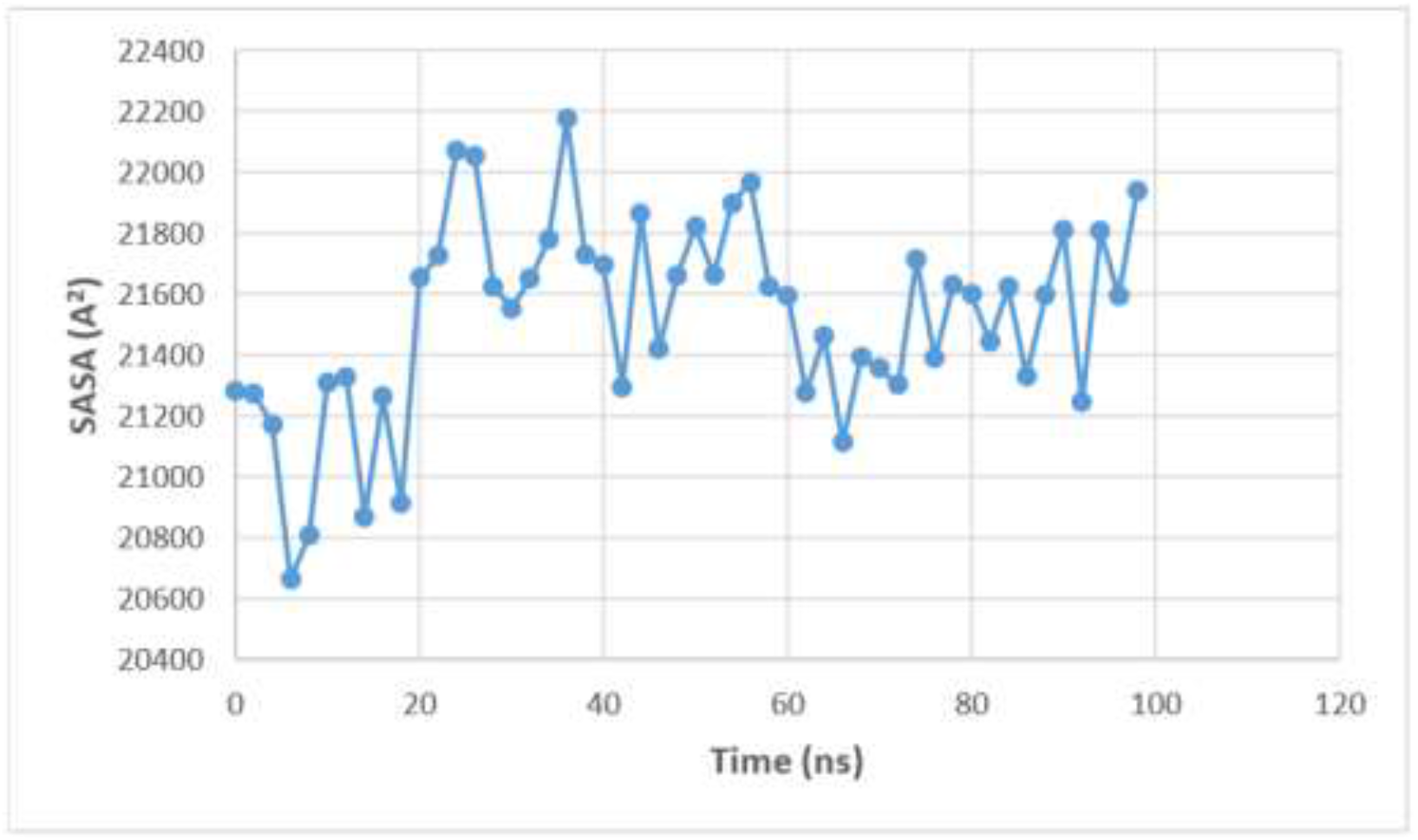
Solvent accessible surface area for the ligand HLA-Cw4-β2m.

Table 4 includes the solvent-accessible surface areas (SASA) for the ligand, receptor and complex. The calculated buried surface area due to binding is about 1559.6 A2, in which it can be calculated by:

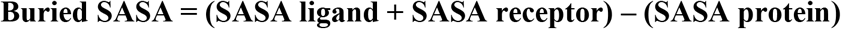

**Table 4.** The solvent-accessible surface areas (SASA) for the ligand, receptor, and complex Values.

| Protein | SASA Avg (Å<sup>2</sup>) | STDV |
| --- | --- | --- |
| HLA-Cw4-β2m-KIR2DL1 complex | 31668.671 | 404.017 |
| Ligand HLA-Cw4-β2m | 21519.433 | 326.181 |
| Receptor KIR2DL1 | 11708.899 | 229.475 |

According to the previous articles, a total of 1,914 Å^2^ of the solvent-accessible surface area is buried, calculated by the surface program for the complex X-ray structure. It is a relatively large buried surface area compared with other peptides–MHC complexes[11].

Figure 13 depicts the final frame of HLA-Cw4-β2m-KIR2DL1 complex structure with small peptide QYDDAVYKL, in which the system contains 348217 atoms (9062 amino acid atoms and 339155 TIP3P water molecules).

**Fig. 12.**
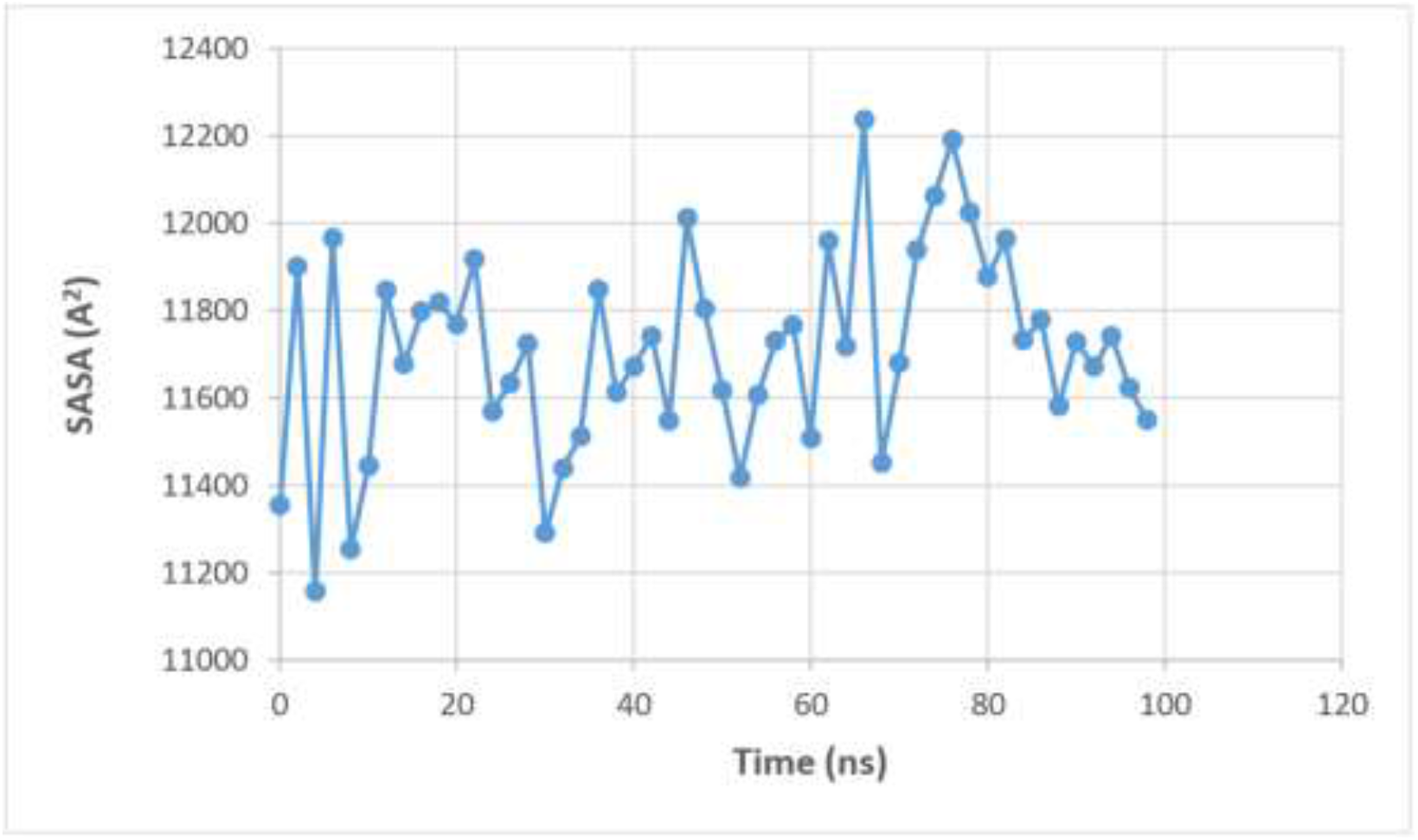
Solvent accessible surface area for the receptor KIR2DL1.

**Fig. 13.**
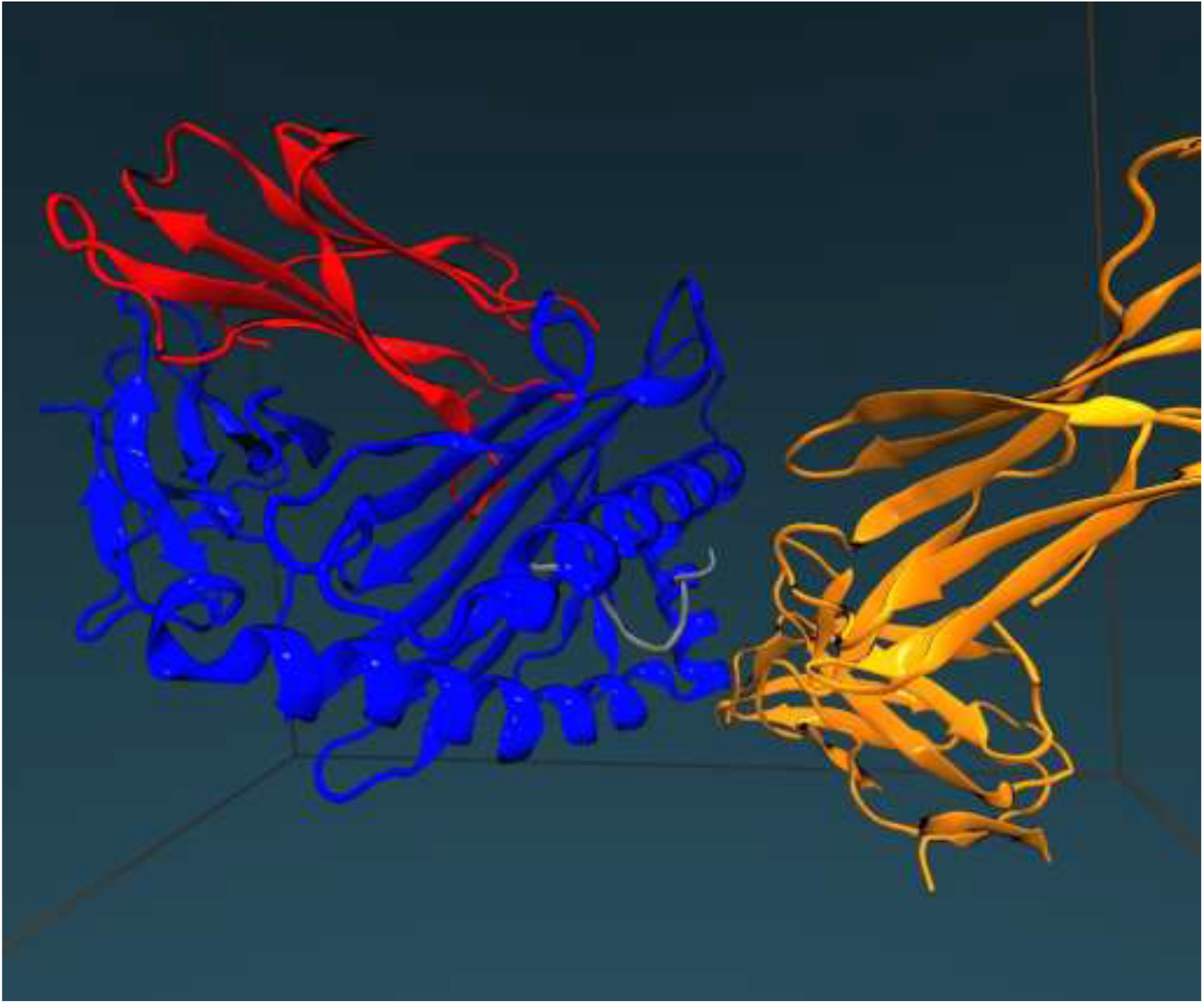
The final structure of HLA-Cw4-β2m-KIR2DL1 protein after 100 *ns*.

**Fig. 14.**
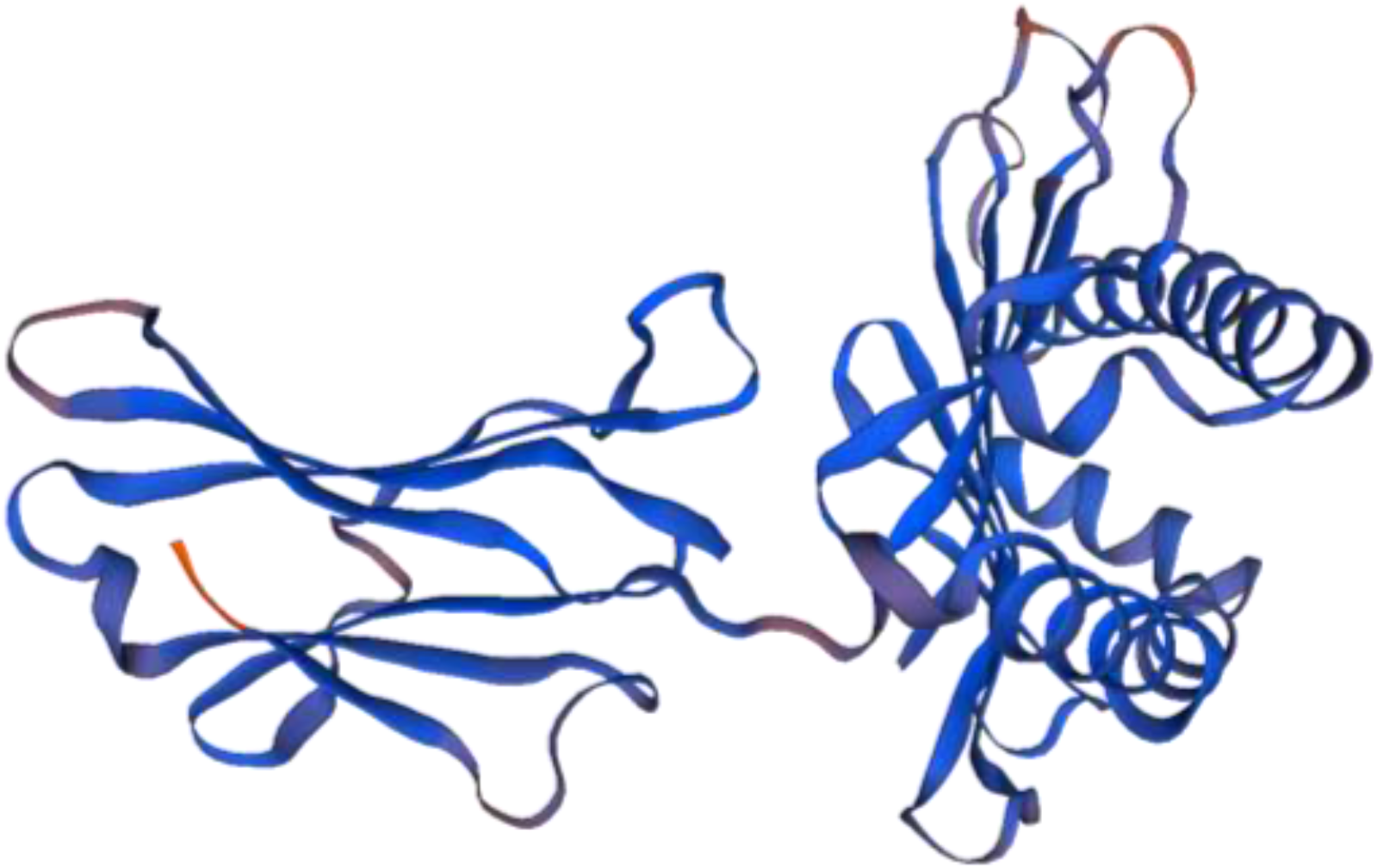
HLA-Cw6-β2m-KIR2DS1 protein obtained using SWISS-MODEL.

### 3.4. Homology Modeling Using SWIS-MODEL

The HLA-Cw6-β2m-KIR2DS1 protein model was built using the SWISS-MODEL program with the selection of segments from HLA structure in the database for matching the segments that was done based on the sequence identity, geometry, and energy. The sequence that was used for the identification of template structures in the PDB (https://www.rcsb.org/)[28] is:

(HSMRYFSTSVSWPGRGEPRFIAVGYVDDTQFVRFDSDAASPRGEPREPWVEQEGPEYWDRETQKYKRQAQADRVNLRKLRGYYNQSEDGSHTLQRMFGCDLGPDGRLLRGYNQFAYDGKDYIALNEDLRSWTAADTAAQITQRKWEAAREAEQRRAYLEGTCVEWLRRYLENGKETLQRAEHPKTHVTHHPVSDHEATLRCWALGFYPAEITLTWQWDGEDQTQDTELVETRPAGDGTFQKWAAVVVPSGEEQRYTCHVQHEGLPEPLTLRWMIQRTPKIQVYSRHPAENGKSNFLNCYVSGFHPSDIEVDLLKNGERIEKVEHSDLSFSGDWSFYLLYYTEFTPTEKDEYACRVNHVTLSQPKIVKWDRDQYDDAVYKLMHEGVHRKPSLLAHPGPLVKSEETVILQCWSDVMFEHFLLHREGMFNDTLRLIGEHHDGVSKANFSISRMTQDLAGTYRCYGSVTHSPYQVSAPSDPLDIVIIGLYEKPSLSAQPGPTVLAGENVTLSCSSRSSYDMYHLSREGEAHERRLPAGPKVNGTFQADFPLGPATHGGTYRCFGSFHDSPYEWSKSSDPLLVSVTGNPSNSWPSPTEPSSKTGNPRHLH)

The best structures were selected according to many factors that determines the accuracy of the structures generated in homology modeling. Such as, Coverage, GMQE, Identity and Oligo State; therefore, according to these variables, we select four eligible templates:

1. Histocompatibility leukocyte antigen HLA-CW4 crystal structure of HLA-CW4, a ligand for the KIR2DL1 natural killer cell inhibitory receptor with values of GMQE =0.39 and Identity = 95.60.
2. HLA class I histocompatibility antigen, B-39 alpha chain an acetylated peptide complexed with HLA-B3901 with values of GMQE =0.42 and Identity = 88.97.
3. HLA-CW3 structure of a complex between the human natural killer cell receptor KIR2DL2 and a class I MHC ligand HLA-CW3 with values of GMQE =0.40 and Identity = 92.70
4. HLA-CW3 structure of a complex between the human natural killer cell receptor KIR2DL2 and a class I MHC ligand HLA-CW3 with values of GMQE =0.40 and Identity = 93.38.

Finally, the PDB file for HLA-Cw6-β2m-KIR2DS1 was constructed using the homology modeling (SWISS-MODEL) based on the HLA-Cw4-β2m-KIR2DL1 protein.

## 4. Conclusions

The results of this molecular dynamics investigation for the complex HLA-Cw4-β2m-KIR2DL1 were performed by NAMD over 100 *ns*, in aqueous solution, reveal the following observations: The molecular dynamics simulation for the complex HLA-Cw4-β2m-KIR2DL1 were performed by NAMD over 100 ns. A simulation time of 100 ns is long enough to achieve equilibrium. Mainly electrostatic forces and minor van der Waals forces contribute to the favorable nonbonded interaction energy observed for antigen (HLA-Cw4) (mostly chain A) interacting with the receptor (KIR2DL1) in water. The interaction of chain C (small peptide) of (HLA-Cw4) does not contribute to the non-bonding interactions between ligand and receptor. Therefore, it does not participate in the receptor (KIR2DL1) specificity to the ligand (HLA-Cw4). The major contribution to the binding site between receptor-ligand comes from Lys^80^ of HLA-Cw4 and some residues of KIR2DL1. Where the side chain of Met44 of KIR2DL1 formed mainly hydrophobic interactions with the side chain of Lys^80^ with high exposure (the Average area of contact is 92.7 A^2^). and Glu^187^ of KIR2DL1 formed mainly large electrostatic interactions with Lys^80^ of HLA-Cw4 (The non-bonding interactions is about −80 kcal/mol). That explains the mutation experimental results, which indicate that mutation in Lys^80^ of HLA-Cw4 causes losing of recognition of the receptor (KIR2DL1) to the ligand (HLA-Cw4).

Many hydrogen bonds formed between the receptor KIR2DL1 and ligand HLA-Cw4, which contribute to specificity of receptor to ligand. For example, H-bond formation between Lys^80^ of HLA-Cw4 and Glu^187^ of KIR2DL1 and Lys^80^ of HLA-Cw4 and Asp^183^ of KIR2DL1. The calculated buried surface area due to binding is about 1559.6 A^2^, which is relatively large compared with other peptides–MHC complexes. Many salt bridges formed between the residues in receptor KIR2DL1 (Chain D) and ligand HLA-Cw4 (Chain A). Remarkably, the salt bridges between Lys^80^ of HLA-Cw4 and Glu^187^ of KIR2DL1 contribute to the specificity of receptor to ligand. Mainly electrostatic forces and minor van der Waals forces contribute to the favorable nonbonded interaction energy observed for antigen (HLA-Cw4) (mostly chain A) interacting with the receptor (KIR2DL1) in water.

## Supporting Information

Snapshots illustrating inter-molecular H-bonds between amino acids from chain D and chain A. The hydrogen bonds between the ligand HLA-Cw4 and its specific peptide and nonbonded interactions between KIR2DL1 and HLA-Cw4.

## Notes

The authors declare no competing financial interests.

## Acknowledgment

Mansour H. Almatarneh is grateful to the Deanship of Academic Research at the University of Jordan for the grant. We also gratefully acknowledge Compute Canada for the computer time.

