## Supporting Information for "Molecular Dynamics Simulations of HLA-CW4-B2M-KIR2DL1 Protein and Homology Modeling of a Complex Associated with Psoriasis Disease (HLA-CW6-B2M-KIR2DS1)"

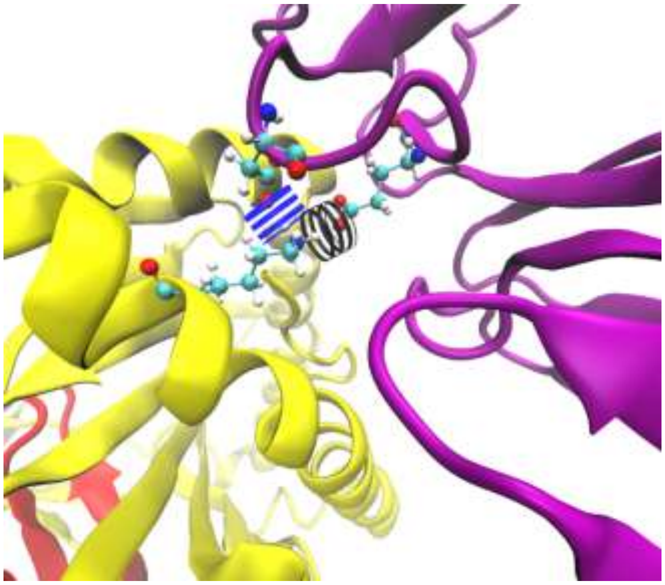

**Fig. S1.** Snapshot illustrating inter-molecular H-bonds between Lys80 from chain A with Asp183 from chain D (the H-bond is indicated by a dashed blue line) and Lys80 from chain A with Glu187 from chain D (the H-bond is indicated by a dashed black line).

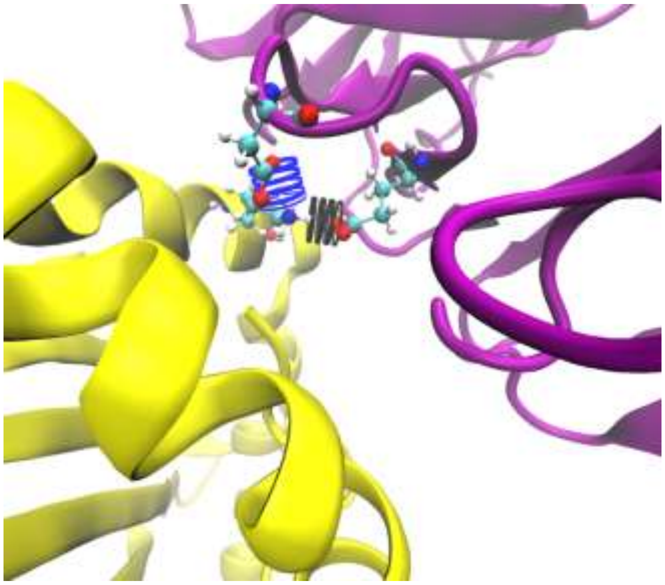

**Fig. S2**. Snapshot illustrating inter-molecular H-bonds between Lys146 from chain A with Asp183 from chain D (the H-bond is indicated by a dashed blue line) and Lys146 from chain A with Glu187 from chain D (the H-bond is indicated by a dashed black line).

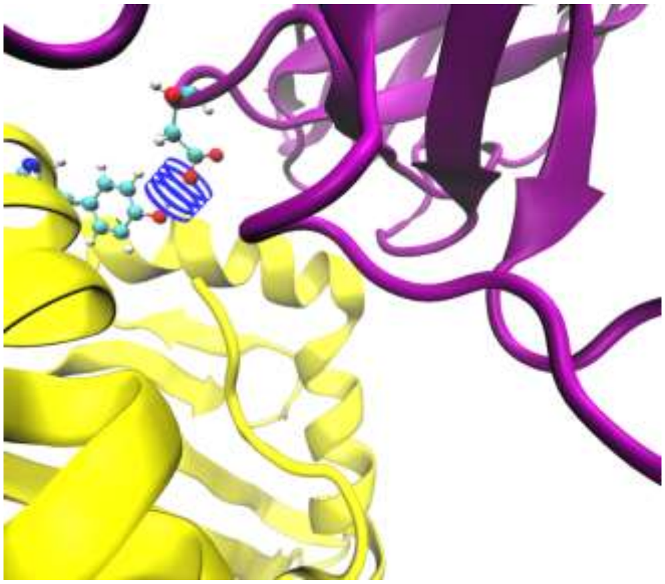

**Fig. S3**. Snapshot illustrating inter-molecular H-bonds between Tyr84 from chain A with Asp183 from chain D (the H-bond is indicated by a dashed blue line).

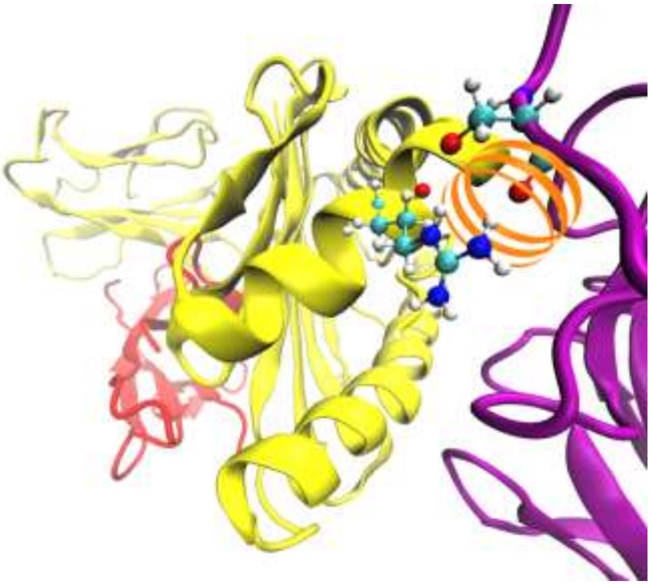

**Fig. S4.** Snapshot illustrating inter-molecular H-bonds between Arg145 from chain A with Ser133 from chain D (the H-bond is indicated by a dashed orange line).

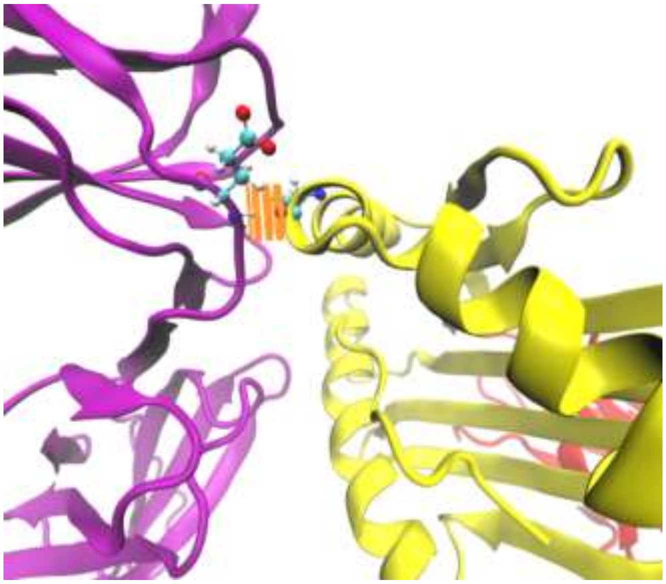

**Fig. S5.** Snapshot illustrating inter-molecular H-bonds between Ala149 from chain A with Glu106 from chain D (the H-bond is indicated by a dashed orange line).

**Table S1**

Hydrogen bonds between the ligand HLA-Cw4 and its specific peptide.

| HLA-Cw4 Specific Peptide | HLA- Cw4 | Occupancy |
|---|---|---|
| PAla$^{5}$-Main-O | Arg$^{156}$-Side-NH1 | 58.00% |
| PLys$^{8}$-Main-O | Trp$^{147}$-Side-NE1 | 56.00% |
| PLeu$^{9}$-Main-N | Asn$^{77}$-Side-OD1 | 50.00% |
| PTyr$^{7}$-Side-OH | Asp$^{74}$-Side-OD1 | 46.00% |
| PAsp$^{3}$-Side-OD1 | Arg$^{97}$-Side-NH1 | 42.00% |
| PAsp$^{4}$-Side-OD1 | Lys$^{66}$-Side-NZ | 42.00% |
| PLeu$^{9}$-Side-OT2 | Lys$^{146}$-Side-NZ | 38.00% |
| PAsp$^{3}$-Side-OD2 | Arg$^{97}$-Side-NH1 | 32.00% |
| PTyr$^{2}$-Main-O | Lys$^{66}$-Side-NZ | 32.00% |
| PGln$^{1}$-Main-O | Tyr$^{159}$-Side-OH | 30.00% |
| PAsp$^{3}$-Side-OD2 | Arg$^{97}$-Side-NH1 | 30.00% |
| PAsp$^{3}$-Side-OD1 | Arg$^{97}$-Side-NH2 | 26.00% |
| PTyr$^{7}$-Main-O | Asn$^{77}$-Side-ND2 | 22.00% |
| PLeu$^{9}$-Side-OT1 | Lys$^{80}$-Side-NZ | 22.00% |
| PGln$^{1}$-Side-NE2 | Glu$^{63}$-Side-OE2 | 18.00% |
| PAsp$^{4}$-Side-OD2 | Lys$^{66}$-Side-NZ | 18.00% |
| PTyr$^{7}$-Main-O | Gln$^{70}$-Side-NE2 | 16.00% |
| PTyr$^{7}$-Side-OH | PGln$^{1}$-Main-N | 16.00% |
| PGln$^{1}$-Main-N | Glu$^{63}$-Side-OE1 | 14.00% |
| PTyr$^{2}$-Side-OH | Gln$^{70}$-Side-OE1 | 14.00% |
| PLys$^{8}$-Side-NZ | Glu$^{152}$-Side-OE1 | 14.00% |

| | | |
|---|---|---|
| PTyr$^{7}$-Main-N | Gln$^{70}$-Side-OE1 | 14.00% |
| PVal$^{6}$-Main-O | Arg$^{156}$-Side-NH2 | 12.00% |
| PAsp$^{3}$-Side-OD1 | Arg$^{156}$-Side-NH1 | 10.00% |
| PTyr$^{7}$-Side-OH | Asp$^{74}$-Side-OD2 | 10.00% |
| PLeu$^{9}$-Side-OT1 | Lys$^{146}$-Side-NZ | 8.00% |
| PGln$^{1}$-Side-NE2 | Tyr$^{159}$-Side-OH | 8.00% |
| PGln$^{1}$-Main-N | Glu$^{63}$-Side-OE2 | 6.00% |
| PTyr$^{2}$-Side-OH | PSer$^{9}$-Side-OG | 6.00% |
| PTyr$^{2}$-Side-OH | Arg$^{97}$-Side-NH1 | 6.00% |
| PLeu$^{9}$-Side-OT2 | Lys$^{80}$-Side-NZ | 6.00% |
| PAsp$^{4}$-Side-OD2 | Tyr$^{159}$-Side-OH | 4.00% |
| PAsp$^{4}$-Side-OD1 | Tyr$^{159}$-Side-OH | 4.00% |
| PTyr$^{7}$-Side-OH | Asn$^{77}$-Side-ND2 | 4.00% |
| PGln$^{1}$-Side-OE1 | Lys$^{66}$-Side-NZ | 2.00% |
| PGln$^{1}$-Side-NE2 | Trp$^{167}$-Side-NE1 | 2.00% |
| PGln$^{1}$-Side-NE2 | Tyr$^{171}$-Side-OH | 2.00% |
| PGln$^{1}$-Side-OE1 | Tyr$^{159}$-Side-OH | 2.00% |
| PAsp$^{3}$-Side-OD2 | Arg$^{156}$-Side-NH1 | 2.00% |
| PAsp$^{3}$-Main-O | Lys$^{66}$-Side-NZ | 2.00% |
| PLys$^{8}$-Side-NZ | Glu$^{152}$-Side-OE2 | 2.00% |

**Table S2**

**Nonbonded Interactions between KIR2DL1 and HLA-Cw4.**

| KIR2DL1 | HLA-Cw4 | Nonbonded | Elec | vdW |
|---|---|---|---|---|
| $Asp^{183}$ | $Lys^{146}$ | -105.681 | -107.447 | 1.7658 |
| $Glu^{106}$ | $Arg^{151}$ | -91.406 | -92.3041 | 0.8981 |
| $Asp^{135}$ | $Arg^{145}$ | -91.0549 | -91.7595 | 0.7056 |
| $Glu^{187}$ | $Lys^{80}$ | -80.0464 | -81.0005 | 0.9542 |
| $Ser^{133}$ | $Arg^{145}$ | -27.4284 | -27.1864 | 0.2421 |
| $Asp^{183}$ | $Tyr^{84}$ | -15.6974 | -16.064 | 0.3666 |
| $Met^{44}$ | $Arg^{79}$ | -8.08945 | -5.3702 | -2.7193 |
| $Tyr^{105}$ | $Lys^{146}$ | -5.5869 | -3.3145 | -2.2724 |
| $Phe^{45}$ | $Arg^{75}$ | -4.2565 | 0.8282 | -4.2565 |
| $Tyr^{105}$ | $Ala^{149}$ | -4.2084 | -2.0087 | -2.1997 |
| $Asp^{72}$ | $Val^{76}$ | -4.1351 | -3.0198 | -1.1153 |
| $Phe1^{81}$ | $Lys^{146}$ | -1.6768 | 1.3306 | -3.0074 |
| $Phe^{45}$ | $Val^{76}$ | -1.2849 | -0.0905 | -1.944 |
| $Ser^{132}$ | $Ala^{149}$ | -1.1198 | -0.3506 | 0.7692 |
| $Phe^{45}$ | $Arg^{79}$ | -0.5779 | 0.009 | -0.587 |
| $Met^{44}$ | $Lys^{80}$ | -0.567 | 2.1283 | -2.6953 |
| $Ser^{184}$ | $Lys^{146}$ | 1.98008 | 2.2654 | -0.2854 |
